# Intron-lariat spliceosomes convert lariats to true circles: implications for intron transposition

**DOI:** 10.1101/2024.03.26.586863

**Authors:** Manuel Ares, Haller Igel, Sol Katzman, John P. Donohue

## Abstract

Rare, full length circular intron RNAs distinct from lariats have been reported in several species, but their biogenesis is not understood. We envision and test a hypothesis for their formation using *Saccharomyces cerevisiae*, documenting full length and novel processed circular RNAs from multiple introns. Evidence implicates a previously undescribed catalytic activity of the intron-lariat spliceosome (ILS) in which the 3’-OH of the lariat tail (with optional trimming and adenylation by the nuclear 3’ processing machinery) attacks the branch, joining the intron 3’ end to the 5’ splice site in a 3’-5’ linked circle. Human U2 and U12 spliceosomes produce analogous full length and processed circles. Post-splicing catalytic activity of the spliceosome may promote intron transposition during eukaryotic genome evolution.

## INTRODUCTION

Circular RNAs are widespread in nature. Among the first discovered were viroids and virusoids, whose replication requires a ribozyme to cleave and ligate monomeric circles from the long linear product of rolling circle replication (Lasda and Parker 2014). Other ribozymes including the group I and group II self-splicing introns produce circular RNAs by performing splicing-related side reactions during or after splicing (Lasda and Parker 2014; Grabowski et al. 1981; Murray et al. 2001). During the normal course of splicing, group II introns and the spliceosome produce tailed circles called lariats closed by a 2’-5’ linkage between the intron 5’ end and a 2’-OH of an internal branchpoint residue (Ruskin et al. 1984; Reed and Maniatis 1985; Konarska et al. 1985; van der Veen et al. 1986; Peebles et al. 1986).

Nucleolytic trimming of lariat 3’ tails can produce tailless lariats: circular intron RNAs closed by a 2’-5’ linkage (Zhang et al. 2013; Gardner et al. 2012; Lasda and Parker 2014; Neil and Fairbrother 2019). Using its standard first and second step splicing chemistry, the spliceosome can also create circular exons through backsplicing, an alternative splicing event that joins a downstream 5’ splice site (ss) to an upstream 3’ss (Nigro et al. 1991; Cocquerelle et al. 1992; Zaphiropoulos 1996; Salzman et al. 2012). If the natural order of the splice sites of group I, group II, or spliceosomal introns are permuted, they can also backsplice to create circular exons (Pasman et al. 1996; Puttaraju and Been 1992; Ford and Ares 1994). Circular introns are also produced by protein-based splicing of tRNA introns (Schmidt et al. 2019), archaeal rRNA and tRNA genes (Lykke-Andersen et al. 1997), and non-spliceosomal introns in protein-coding mRNAs such as for *S. cerevisiae HAC1* (Schmidt et al. 2019; Cherry et al. 2019) or numerous genes in *Euglena* (Gumińska et al. 2021). In addition to providing important insight into splicing mechanisms, some circular RNAs are functional (Lasda and Parker 2014; Neil and Fairbrother 2019), although the biological roles for most remain mysterious.

During their analysis of RNAseq reads to identify intron lariat branch points, Fairbrother and colleagues noted that ∼3% of their intron-derived split reads appeared to show "branch formation" at the 3’ss (Taggart et al. 2017, 2012). They validated the existence of intron-derived RNAs in which the 3’ss is directly joined to the 5’ss, but noted the reads lacked characteristic mutations created by reverse transcriptase (RT) reading through the lariat branch, suggesting they were not 2’-5’ linked, and thus not branch points. They speculated that the circles were formed after splicing by an unknown ligation mechanism. Similar circles have since been documented in the distantly related eukaryotic microbe *Entamoeba histolytica* (Mendoza-Figueroa et al. 2018). To distinguish them from tail-trimmed lariats that have also been called intron circles (Zhang et al. 2013; Gardner et al. 2012; Lasda and Parker 2014; Neil and Fairbrother 2019), we refer to this class as "true circles". Group II introns can also form circles that join their 5’ss to their 3’ss through a 2’-5’ linkage (Murray et al. 2001). We wondered whether the spliceosome creates intron circles and if so how. We searched in *Saccharomyces cerevisiae*, and found full length intron circles for multiple introns, along with a new class of processed intron circles. We provide evidence supporting a model for their formation that implicates persistent catalytic activity of the spliceosome after spliced exon release. By analogy to the transposable group II introns, we suggest that the modern spliceosome may still be able to transpose introns to new locations.

## RESULTS

### How could the spliceosome create true intron circles?

In addition to its two forward splicing reactions, branch formation (step 1) and exon ligation (step 2), the spliceosome catalyzes both reverse reactions (Tseng and Cheng 2008). In the reverse of the first step, exon 1 (E1) attacks the branch phosphate to regenerate the pre-mRNA. If the lariat tail could attack the branch phosphate in a similar way, a circular intron would be formed (Fig 1A, top). We hypothesize that after the spliced exons are removed from the spliceosome, the resulting intron-lariat spliceosome (ILS) would have an empty E1 binding site that could accommodate the lariat tail. Remodeling of the ILS into a C-complex like conformation with the branch phosphate in the catalytic center could allow the lariat tail 3’-OH to attack and create a circular intron (Fig 1A, bottom). To explore this hypothesis in a tractable system we turned to yeast.

**Figure 1.**
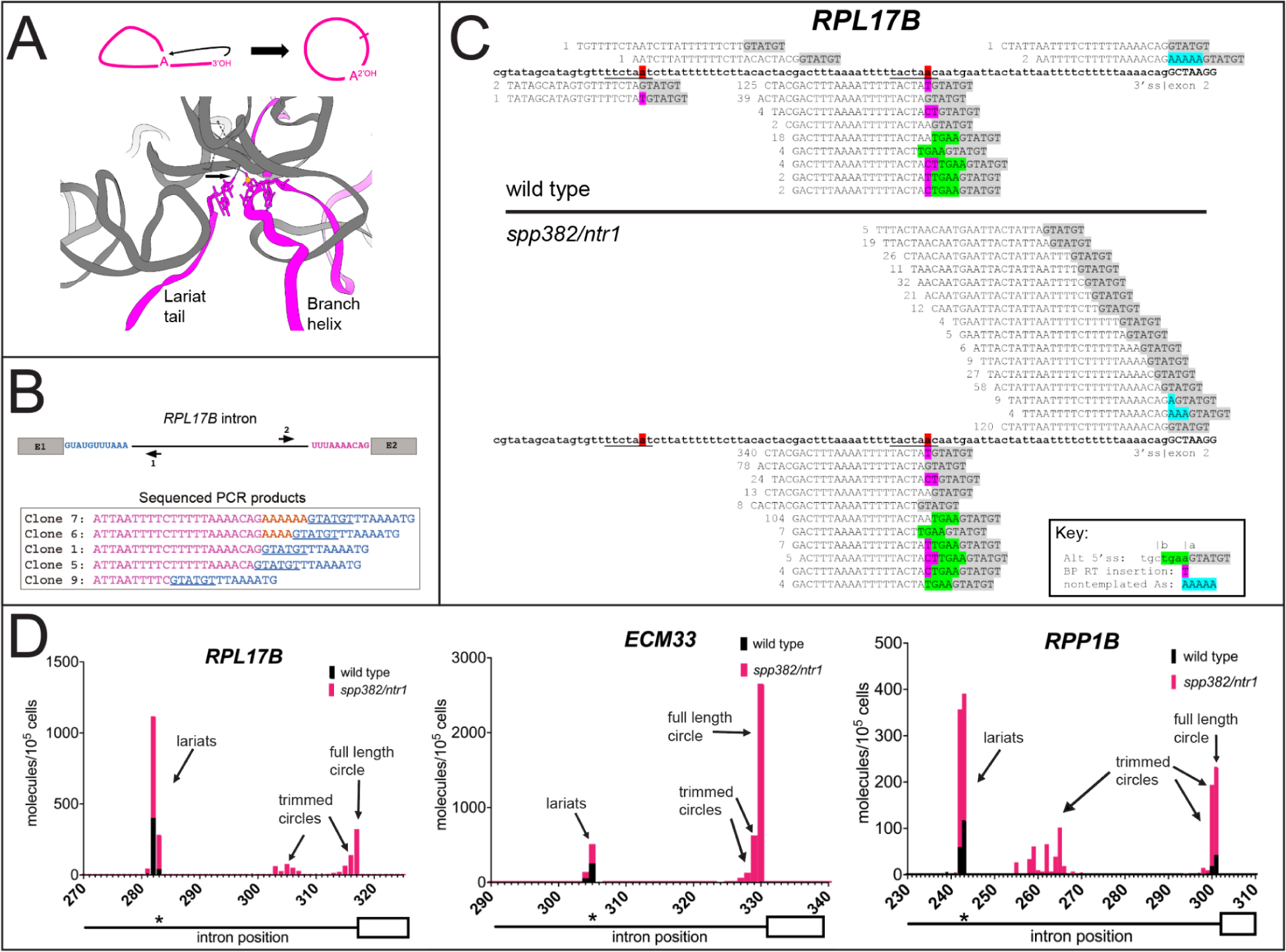
A hypothesis, and detection of intron circles in *S. cerevisiae*. (**A**) A possible mechanism for the spliceosome to create intron circles by nucleophilic attack of the lariat tail 3’-OH (arrow, top and bottom) on the branch phosphate (yellow sphere, bottom), forming a 3’-5’ junction between the 3’ss and the 5’ss (top). A remodeled intron-lariat spliceosome (ILS) in which the lariat tail 3’-OH is bound in E1 binding site (bottom, arrow) could carry out a reaction formally similar to the reverse of step 1 (reverse of branch formation) to create the intron circle (bottom). Cartoon derived from C-complex model 5GMK (Wan et al. 2016), using ChimeraX (Meng et al. 2023). Only RNA is shown, U2, U6, and U5 in gray, intron in magenta. (**B)** *RPL17B* intron with "inverse" primers to capture circular intron junctions. Sequences of selected cloned PCR products are shown, with the 5’ss underlined and non-genomic As in gold. (**C)** High throughput sequencing reveals a complex set of processed circular *RPL17B* introns whose abundance is dramatically increased by the *spp382-1* mutation. The intron is in bold. Reads are aligned above (circles) or below (lariats), the 5’ss sequence highlighted in gray, and non-genomic As in blue. Deduplicated (unique) read counts from each library are shown in front of each read. Lariats derived from incorrect 5’ss are in green. (**D)** Different introns produce distinct distributions of intron circles. Graphs show the junction locations (x-axis) and number of unique reads with junctions at that location (y-axis, calibrated to a spiked-in circular RNA) per 10^5^ yeast cells, from wild type (black bars) or *spp382-1* (pink bars) cells. The 3’ part of each intron (line) and its second exon (white box) are shown below the x-axis, with the asterisk indicating the position of the bp. Additional introns and data are in Fig S1 and Table S1.

### Full length and novel "processed" circular intron RNAs from multiple *S. cerevisiae* genes

To search for intron circles, we designed outward pointing ("inverse") PCR primers for a highly expressed intron-containing gene *RPL17B*, and used them to amplify, clone, and sequence reverse transcripts extending from the 5’ end of the intron past predicted circle junctions (Fig 1B, Table S1). In addition to RNAs containing junctions between the 5’ss and 3’ss of the *RPL17B* intron analogous to those from mammals and *Entamoeba* (clone 1), we find unexpected circles that include nontemplated adenosine residues (As) between their 5’ and 3’ss (clones 6 and 7), as well as those missing nucleotides from the 3’ end of the intron (clones 5 and 9). To test other introns, as well as to sample physiological, biochemical, and genetic conditions that might reveal insights into intron circle formation, we devised a multiplexed circular intron amplicon library preparation protocol using inverse primers carrying unique molecular indexes (UMIs), and sample barcoded adaptors for Illumina sequencing of a dozen introns (See Supplemental Methods). We sequenced libraries from wild type and a mutant strain (*spp382-1)* defective in spliceosome disassembly (Pandit et al. 2006), and readily detected circles for all 12 tested introns. We examined the reads from the *RPL17B* (Fig 1C) and *ACT1* (Fig S1) introns in detail. For *RPL17B* in wild type cells (Fig 1C, top), lariats (5’ss joined to the bp) are readily observed, at both the annotated (TACTA<u>A</u>C) and an upstream cryptic bp (TTCTA<u>A</u>T). The RT mutational signature at the bp is evident (in purple) as is the use of an incorrect 5’ss upstream (green). Reads derived from an unprocessed full length circular intron (like clone 1 in Fig 1B) as well as from a circular intron with 5 extra non-genomic A residues (like clone 7) are observed. Two circles mapping upstream of the annotated *RPL17B* bp from the wild type library would seem to rule out the hypothesis that circles form when the 3’ end of the lariat tail attacks the branch, since they appear to have been trimmed well past the annotated bp. Note however that they map downstream of the cryptic bp, and could be derived from lariats branched at the cryptic bp via the proposed mechanism. In the *spp382-1* mutant (Fig 1C, bottom), a more complex and numerous group of circles missing nested sets of "tail" nucleotides is observed. We conclude that like humans, rodents, and *Entamoeba* (Taggart et al. 2017, 2012; Mendoza-Figueroa et al. 2018), *S. cerevisiae* makes full length intron circles, and in addition a novel class of processed intron circles. This processed class of circles can also be formed by the proposed spliceosomal mechanism if the lariat tail retains a 3’-OH after processing.

To estimate the numbers of circles per cell, we prepared libraries from RNA samples spiked with known numbers of a synthetic circular RNA molecule for normalization (see Supplemental Methods). We find that intron circles are rare, occuring at ∼1-10 per intron per 10,000 wild type cells (Fig 1D, Fig S1). The same classes of intron RNA circles observed for the *RPL17B* (Fig 1C) and *ACT1* introns (Fig S1) are also found (in different proportions) for other introns. In the *spp382-1* mutant, circles increase to different extents for different introns: not at all for *RPL24B*, and ∼10-100s fold for others (Fig 1D, Fig S1, also see below). In addition some introns produce abundant processed circles (e. g. *RPL17B*, *ACT1, RPP1B*), whereas others (e. g. *RPL19A*, *ARF2*) produce only untrimmed circles (Fig 1D, Fig S1). Taken together these initial observations show that circular intron RNAs arise by a rarely occurring but general process that operates on most or all introns, and that idiosyncratic differences in transcription rate and intron structure likely contribute to the different numbers and types of circles produced from each intron. Increased circular introns from the *spp382-1* mutant provides the first hint that circles are formed by a reaction that is in competition with catalytic inactivation of the spliceosome. Furthermore the processed intron circles fit the profile expected for RNAs that have visited the nuclear exosome and the TRAMP complex (Chlebowski et al. 2013; Zinder and Lima 2017) before they become circular, suggesting that if they are formed by the spliceosome, the lariat tail can exit and reenter before circularization.

### Intron circles are associated with the spliceosome and are not formed by tRNA ligase

Our working hypothesis invokes the spliceosome as the enzyme that closes the intron circles. The only other known RNA ligase in yeast is Trl1/Rlg1 (Phizicky et al. 1992), which ligates tRNA and *HAC1* exons together during non-spliceosomal (tRNA) splicing in yeast (Cherry et al. 2018, 2019; Phizicky et al. 1992; Peschek and Walter 2019). We asked whether circular introns are associated with the spliceosome *in vivo*, and separately whether their formation depends on Trl1 (Fig 2). We tagged the spliceosomal protein Cef1 at its C-terminus with a protein A (TAPS) tag in the *spp382-1* mutant and performed IgG-Sepharose chromatography to capture spliceosomes, detecting the presence of intron circles from *ECM33* in the fractions (Fig 2A, B). To follow the spliceosome through the purification we used ^32^P-labeled oligonucleotides complementary to U2 snRNA (as a spliceosome marker) and SCR1 (a cytoplasmic SRP RNA) for detection by primer extension with RT (Fig S2). The *ECM33* intron circle is clearly enriched in the bound (lane P) spliceosomal fraction of the Cef1-tagged extract but not in the untagged extract (compare lanes 3 and 6, Fig 2B), supporting the idea that the spliceosome forms the circular introns RNAs. To address the role of the tRNA ligase, we compared counts of circular introns (normalized to the spike-in control circle) in wild type and a strain lacking the *TRL1* gene (Cherry et al. 2018), and find that circle formation remains robust in the absence of the tRNA ligase Trl1 (Fig 2C). Based on these observations we conclude that circular introns are most likely created by an RNA ligation activity of the spliceosome.

**Figure 2.**
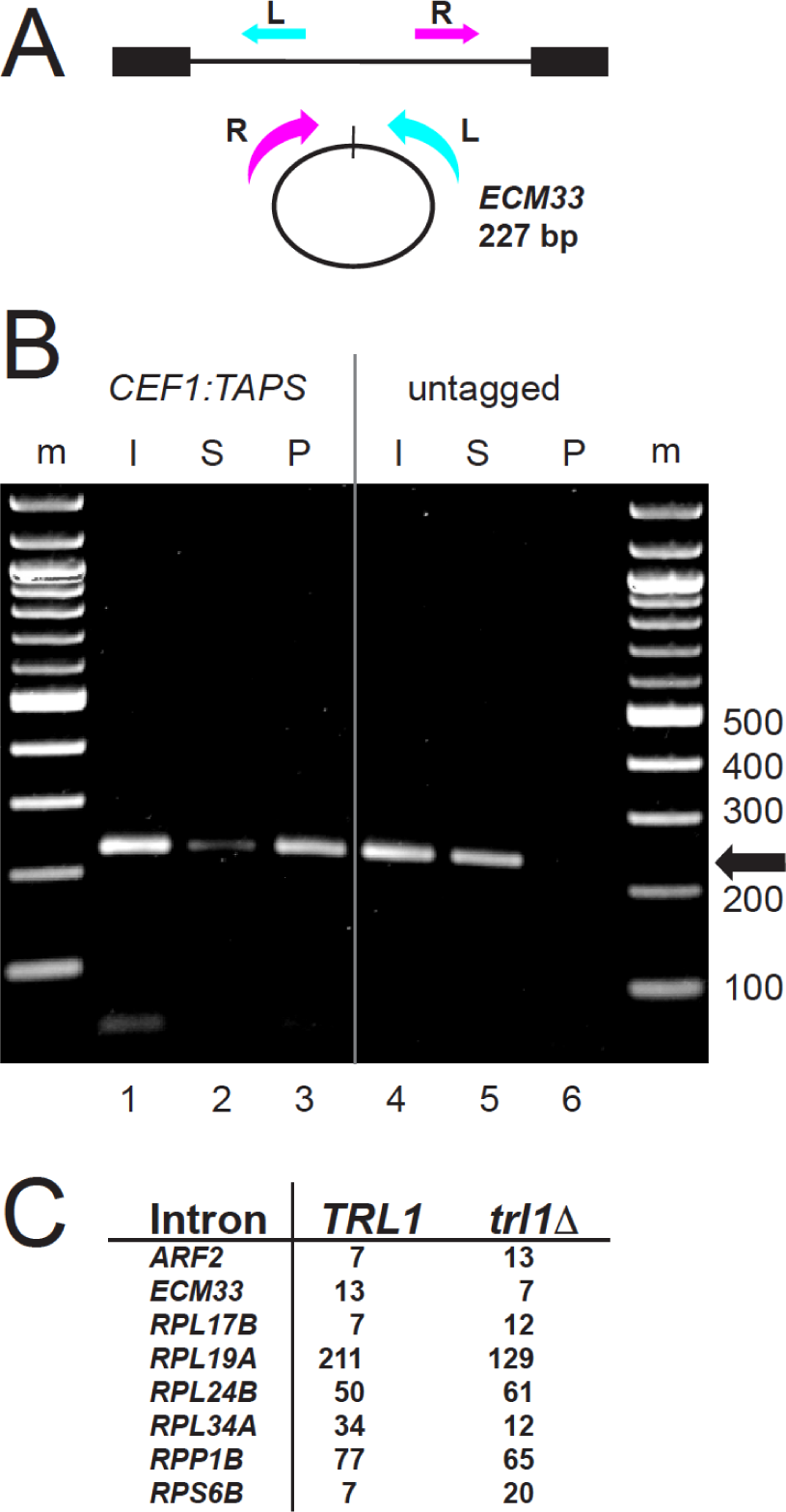
Intron circles are associated with the spliceosome and are not formed by the tRNA ligase Trl1. (**A**) Detecting circular *ECM33* introns. Inverse PCR primers that create a 227 bp product from full length *ECM33* intron circles are shown; the R primer is 3’ of the branch point and cannot detect lariats. (**B**) Association of *ECM33* intron circles with the spliceosome in *spp382-1* cells. The *spp382-1* mutant was fitted with a TAPS tag at the C-terminus of its *CEF1* coding region for affinity purification of spliceosomes. Extracts from this strain and an untagged control were prepared (Input, I) and bound in batch to IgG-sepharose. Unbound (Supernatant, S) and bound (Pellet, P) fractions were prepared. RNA from each fraction was used for RT-PCR to identify *ECM33* circles (227 bp product). By primer extension the pellet fraction from the tagged (but not the untagged) extract is enriched in spliceosomal U2 snRNA and free of cytoplasmic SCR1 (Fig S2). (**C**) Robust detection of intron circles in a strain lacking tRNA ligase. The table shows spike-in normalized circular intron counts for 8 introns from libraries prepared from strains that differ by deletion of *TRL1.* Detailed data are in Table S2.

### Intron circles are not closed by a 2’-5’ linkage

Intron circles could be formed by one of the known activities of the spliceosome (either 2’-5’ or 3’-5’ linkage forming), or by some other activity. The absence of a mutational signature at the circle junction (Taggart et al. 2017, Fig 1C) might not permit the conclusion that intron circles are closed by a 3’-5’ linkage if an internal (i. e. unbranched) 2’-5’ linkage is being formed and does not similarly impede RT, for example as for a group II intron (Murray et al. 2001). To resolve this directly we sought to distinguish lariats (with 2’-5’ branched junctions) and circles with at least one internal non-branched 2’-5’ linkage from true circles (containing only 3’-5’ linkages). The lariat debranching enzyme Dbr1 hydrolyzes 2’-5’ linkages if they are part of a branched structure (Montemayor et al. 2014; Clark et al. 2022), but we were unable to find published evidence of its ability to cleave internal, unbranched 2’-5’ linkages. We designed several synthetic RNA oligonucleotides that differ by replacement of a single 3’-5’ linkage with unbranched 2’-5’ linkage and treated them with purified recombinant yeast Dbr1 protein (Khalid et al. 2005) as described (Qin et al. 2016, kind gift of Aiswarya Krishnamohan and Jon Staley, Fig 3A). Dbr1 cleaves the oligos containing internal 2’-5’ linkages (lanes 2 and 6), without substantially cleaving 3’-5’ linkages (lanes 8 and 10), and therefore can be used to distinguish the classes of molecules described above. We prepared libraries from equivalent amounts of *spp382-1* RNA either treated with Dbr1 or not, and sequenced them, tabulating the numbers of lariats and circles for several introns (Fig 3B). As expected, Dbr1 treatment greatly reduces the recovery of reads corresponding to lariat RNAs, however intron circles in the same digest are resistant. We conclude that full length and processed intron circles are not closed by either a branched or internal 2’-5’ linkage, and are therefore likely to be composed entirely of 3’-5’ linkages. The bond formed by a reaction similar to the reverse of step 1 would be 3’-5’ linked, consistent with the hypothesis that circles are formed as proposed (Fig 1A).

**Figure 3.**
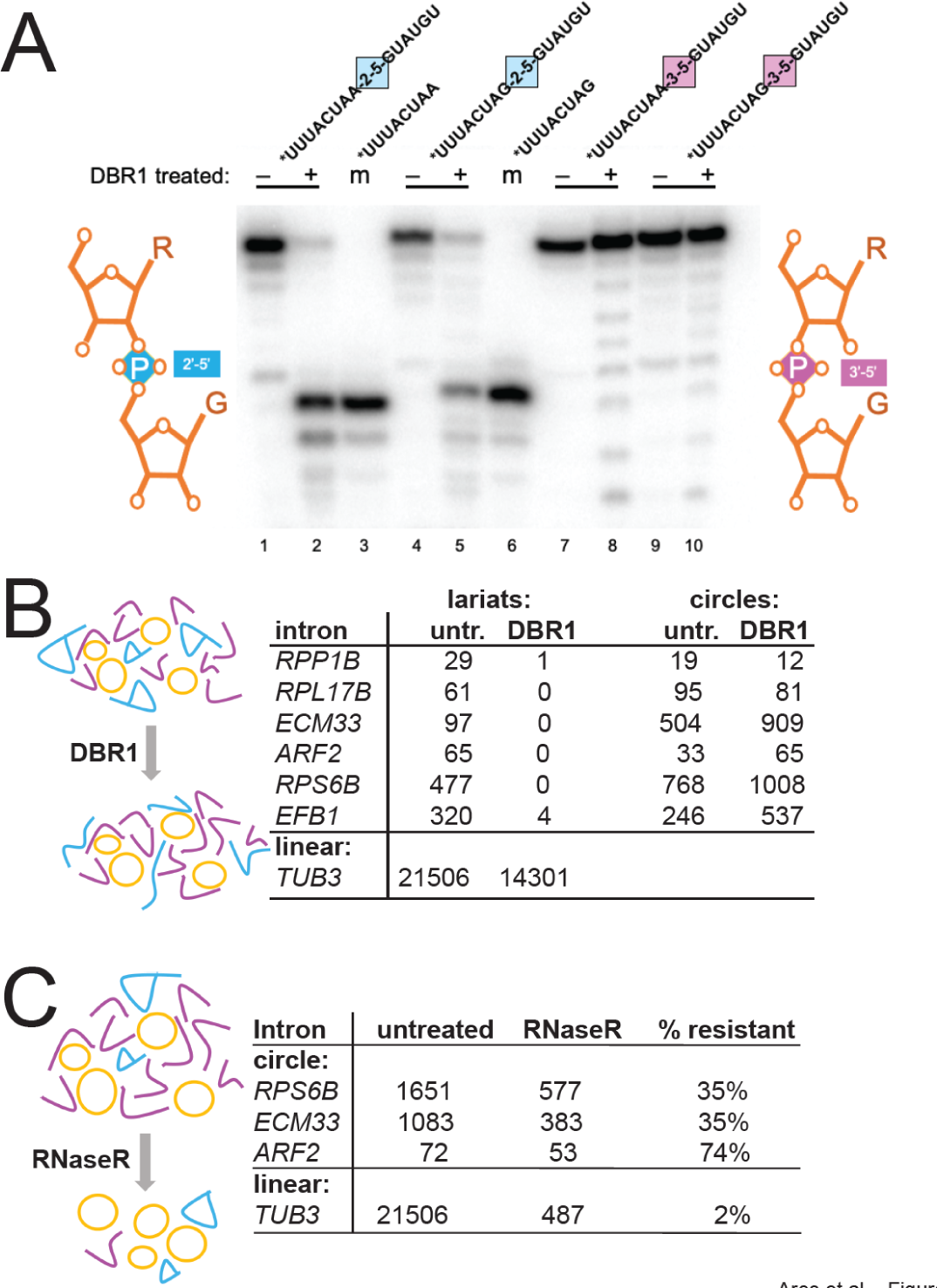
Enzymatic tests of circular intron structure. (**A**) Dbr1 does not require a 3’-nucleotide at the branch and can cleave an internal 2’-5’ linkage. ^32^P-5’ end-labeled RNA oligonucleotides carrying either a single 2’-5’ internal linkage in the middle of an otherwise 3’-5’ linked RNA, or carrying only 3’-5’ linked nucleotides were incubated (or not) with recombinant DBR1 and separated by electrophoresis on an 8M urea, 20% acrylamide gel. Two different substrate sequences (bpA-G5ss and 3ssG-G5ss) were made with either a 2’-5’ (blue) or 3’-5’ (pink) linkage at the junction marked by the colored square are shown above each reaction. Minus and plus signs indicate enzyme addition. Two 8-mers identical to the expected Dbr1 cleavage products were labeled as markers (m, lanes 3 and 6). Comigration with the marker indicates cleavage at the 2’=5’ linkage. (**B)** Circle junctions from the *spp382-1* mutant are resistant to cleavage with Dbr1, whereas lariat junctions are not. Read counts of lariats and circles for six introns from libraries made with or without prior treatment of the input RNA with Dbr1. (**C**) Circle junction-containing RNAs from the *spp382-1* mutant are much more resistant to RNAseR than a linear pre-mRNA. Read counts of circles for three introns from libraries made with or without prior treatment of the input RNA with RNAseR. Detailed data are in Table S3.

### Intron RNAs containing putative circle junctions are resistant to RNAseR

RNA sequencing captures cDNA sequences from junctions consistent with circles, but does not prove the original RNA templates were circular. We tested the resistance of the putatively circular RNAs to the strict 3’ exonuclease RNAseR (Vincent and Deutscher 2006, Fig 3C). Equivalent amounts of *spp382-1* RNA were either treated with RNAseR or not, and sequenced. More than 30% of reads with circle junctions are resistant to RNAseR as shown for the three most abundant intron circles (Fig 3C). In contrast, the recovery of reads for the linear *TUB3* control pre-mRNA was only 2%. Some circles may have broken in the cell or during isolation, and thus were lost to RNAseR, however intron RNAs carrying circle-closing junctions are at least ∼15-30 times more resistant to RNAseR than the linear control intron, and therefore very likely circular.

### Mutations in splicing termination factors promote accumulation of intron circles

The general increase in intron circles per cell in the *spp382-1* mutant is consistent with the idea that delaying spliceosome disassembly (Pandit et al. 2006) promotes circle formation (Fig 1C, D, Fig S1). To explore this further we compared numbers of intron circles (normalized to the spiked in circle) from wild type and strains carrying mutations that slow catalytic inactivation and disassembly of the spliceosome, Prp43 (Toroney et al. 2019; Fourmann et al. 2017; Su et al. 2018; Arenas and Abelson 1997; Martin et al. 2002; Boon et al. 2006; Tsai et al. 2005) and U6 snRNA (Toroney et al. 2019), using Fisher’s exact test (Fig 4). The log odds (fold change) and p values derived using Fisher’s exact text for each intron in each of the three mutants are shown in the inset tables adjacent to each factor’s location in a model of the ILS poised for catalytic inactivation (Wan et al. 2017, 5Y88, Fig 4). For most introns, inhibiting disassembly promotes circle formation (Fig 4), consistent with the conclusion that circle formation is promoted by delaying spliceosome disassembly. Interestingly, the impact on accumulation of individual introns is different for the three mutations. For example, *RPL24B* intron circle formation is unaffected by partial loss of function of any of the termination factors, whereas *ECM33* circle formation is increased ∼250-fold by *spp382-1*, 14-fold by shortening of the U6 3’ end by 5 nucleotides, and 2.5-fold by the *prp43* cold sensitive mutation Q423N (Fig 4, Toroney et al. 2019). The differences between mutants acting on the same intron could be due to qualitative effects of loss of function or quantitative differences in the severity of the tested allele, or both. For example, the greater magnitude of effect of *spp382-1* could be explained if Spp382/Ntr1 binding inhibits catalytic activity before Prp43 function. Differences between introns for the same mutant may be due to differences in the intron-specific intrinsic rate(s) of disassembly steps for spliceosomes carrying different introns. We conclude that the rate of circle formation or the stabilization of intron circles is enhanced when binding of Spp382/Ntr1 and catalytic inactivation of the ILS is delayed, consistent with the idea that circles are formed by the ILS.

**Figure 4.**
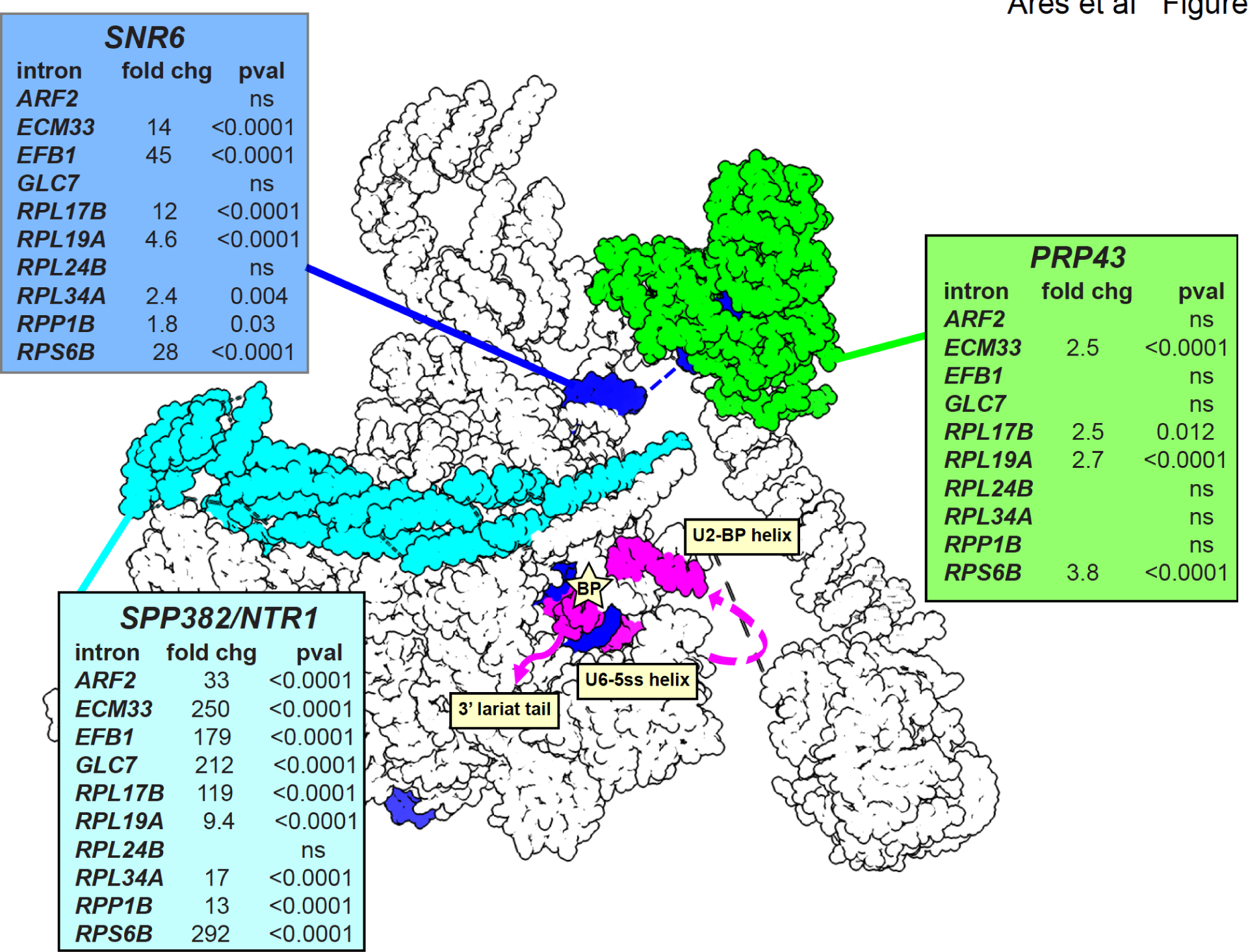
Effects of spliceosome disassembly mutants on abundance of different intron circles. Catalytic inactivation of the spliceosome is mediated by the Prp43 helicase (green) acting in concert with its G-patch protein Spp382/Ntr1 (cyan) on the 3’ end of U6 snRNA (*SNR6*, blue) to pull U6 from the spliceosome (Toroney et al. 2019). Intron circles were counted in pairs of strains carrying wild type or mutant alleles of each factor, normalized using a spiked in circle and tested for significant change using Fisher’s exact test (detailed data are in Table S4). Each table shows the log odds calculation as fold change and the two-sided p value for each of 10 intron comparisons. ns, not significant. The position of the intron is shown in magenta. Image made using ChimeraX (Meng et al. 2023) and the *S. cerevisiae* ILS structure model (5Y88, (Wan et al. 2017).

### Effect of nuclear 3’ decay factors on circle formation

We observe a novel class of intron circles whose 3’ tails have been processed prior to the circularization reaction (Fig 1). Nuclear 3’ RNA processing machinery includes the nuclear exosome with its exonuclease subunits Rrp6 and Dis3, and the TRAMP complex, with either of its two paralogous polyA polymerases Trf4 or Trf5 (Chlebowski et al. 2013; Zinder and Lima 2017). RNA 3’ ends are trimmed and polyadenylated using combinations of these activities in a quasi-distributive fashion coordinated by the Mtr4 RNA helicase (Chlebowski et al. 2013; Zinder and Lima 2017). To test whether circle formation might be influenced by loss of the nuclear 3’ processing machinery we made libraries from deletion strains lacking Rrp6 or Trf4 and compared the yield of different kinds of circles. Three introns (*RPL19A*, *RPP1B*, and *RPL24B*) produced enough circles for statistical analysis (Table S5). Although we could not detect an effect of loss of Trf4 (possibly because its paralog Trf5 remains active in these strains), loss of Rrp6 significantly reduces the number of intron circles from *RPL24B* (∼2 fold, p<0.001 Fisher’s exact test, Table S5) and *RPP1B* (∼3.7 fold, p<0.0001, Fisher’s exact test, Table S5). In contrast, the number of intron circles from *RPL19A* is unaffected (Table S5). Whereas most of the circles from *RPL24B* and *RPP1B* are processed, *RPL19A* produces almost entirely unprocessed circles. A feature that distinguishes these introns is their bp–3’ss distance: *RPL24B* and *RPP1B* have long tails (58 and 65 nt, respectively) and RPL19A has a short tail (14 nt). Taken together, this suggests that introns differ in their interactions with the exosome based on the length of their tails, and that Rrp6 function may enhance the efficiency of circle formation for introns with long tails.

To evaluate this idea in more detail we analyzed the more numerous trimmed circles from the *spp382-1* mutant, comparing their abundance to the length of tail remaining after processing for 10 introns with a range of tail lengths (Fig 5B, see also Fig 1C, Fig S1). Two things are evident, one is that introns with tail lengths >22 can produce trimmed circles that contain as few as 15-20 nucleotides downstream of the branch, but not fewer. Secondly, two introns with tails ≤17 nt (*RPL19A* and *ARF2*) form robust numbers of circles virtually all of which are untrimmed (Fig 5B), although a small number have non-genomic As. This suggests that full length circle formation can occur for introns with at least 14 bp–3’ss nucleotides (and possibly fewer), but that trimming requires introns to have longer bp–3’ss tails. The 15-20 nt tail limit below which trimmed circles are not observed is likely enforced by the distance between the bp A and the closest approach of the exosome active site on the outside of the spliceosome. The observation of non-genomic As but not trimming in the *ARF2* and *RPL19A* intron circles indicates that the TRAMP complex can react with shorter tails than the exosome, possibly because its active site is nearer to its surface and the surface of the spliceosome. We conclude that the nuclear 3’ processing machinery interacts with the set of introns with bp–3’ss distances ("tails") longer than about 17 and enhances circle formation for some of these by shortening the lariat tail, possibly increasing the ease with which it can return to the spliceosome and access the E1 binding site for circularization.

**Figure 5.**
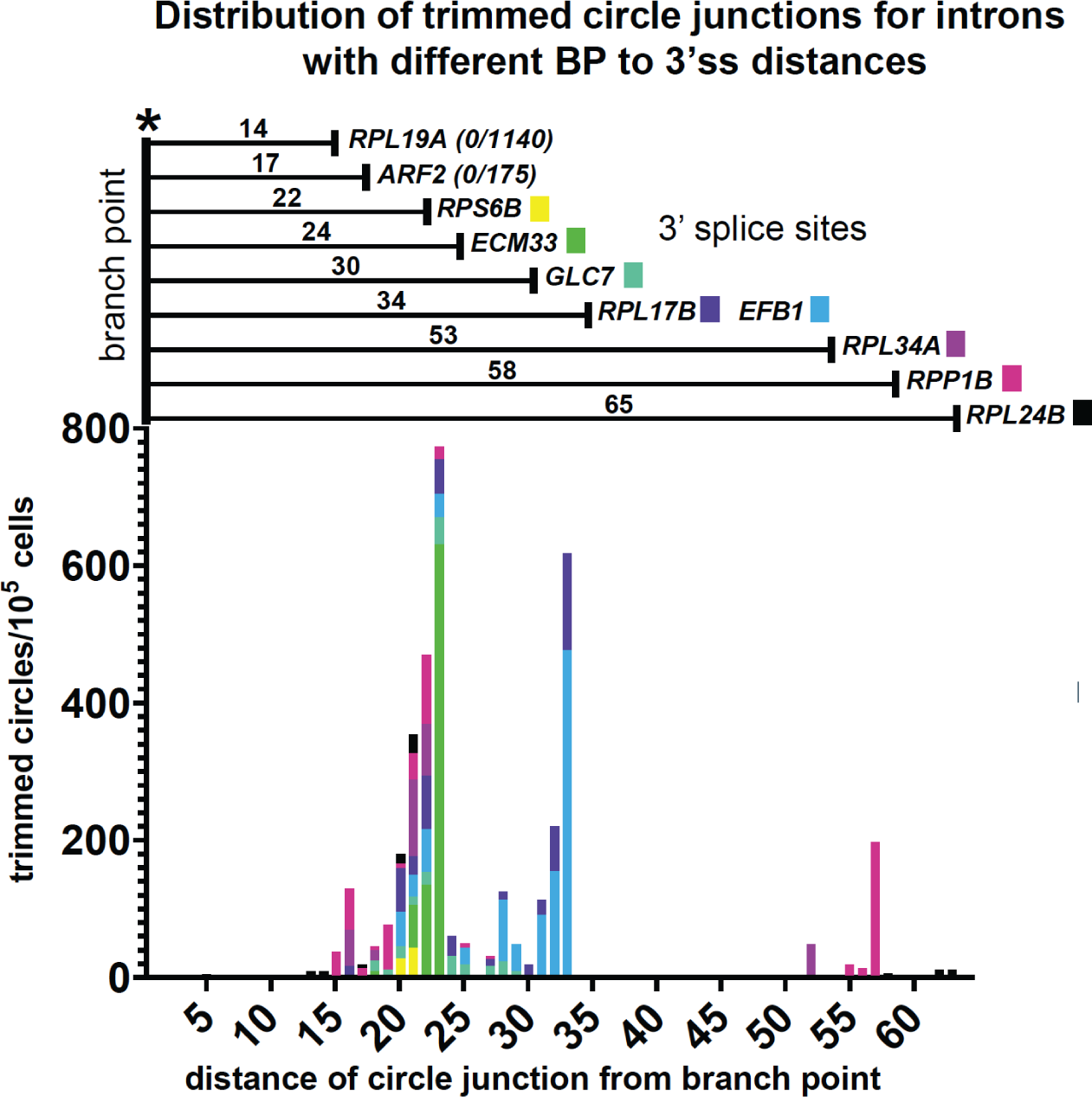
Effects of nuclear 3’ processing on abundance and structure of different intron circles. (**A**) The distribution of trimmed intron circles for all tested introns has a limit of 15-20 nt of lariat tail remaining, independent of the initial tail length. Two introns with natural BP-3’ss tails of 14 (*RPL19A*) or 17 (*ARF2*) nt are not trimmed. Detailed data are in Table S5.

### Human U2 and U12 spliceosomes produce analogous full length and processed intron circles

In their analysis of human and rodent intron circles, Fairbrother and colleagues (Taggart et al. 2017, 2012) describe introns joined by the last intron nucleotide for the U2 major spliceosome, but did not note the presence of circles among the minor (U12) class of introns. They also did not report processed circles, possibly because these can easily be missed when mapping split reads to the genome, especially when non-templated nucleotides are added. To ask whether human intron circles are processed, we analyzed human branchpoint mapping data from a different group (Mercer et al. 2015) by creating a 100 nt permuted intron target file in which the 50 3’ nts of each annotated intron is joined to its 50 5’ nts, for each annotated intron in the human genome (see Methods), and mapped the reads to this target (Fig 6, Table S6). Examples of unprocessed full length intron circles and circles missing 3’ nucleotides, with or without non-genomic As, were readily detected for introns processed by both the major and minor spliceosomes (Fig 6). In addition, using a similar mapping strategy for yeast, we found circular intron reads for 32 of the ∼270 expressed introns (including 7 of the 12 amplicon targets) in standard RNAseq libraries made from rRNA-depleted total RNA (Table S6). Considering previous observations, we suggest that intron circularization can occur for many different introns, in many eukaryotes, for both major and minor class introns.

**Figure 6.**
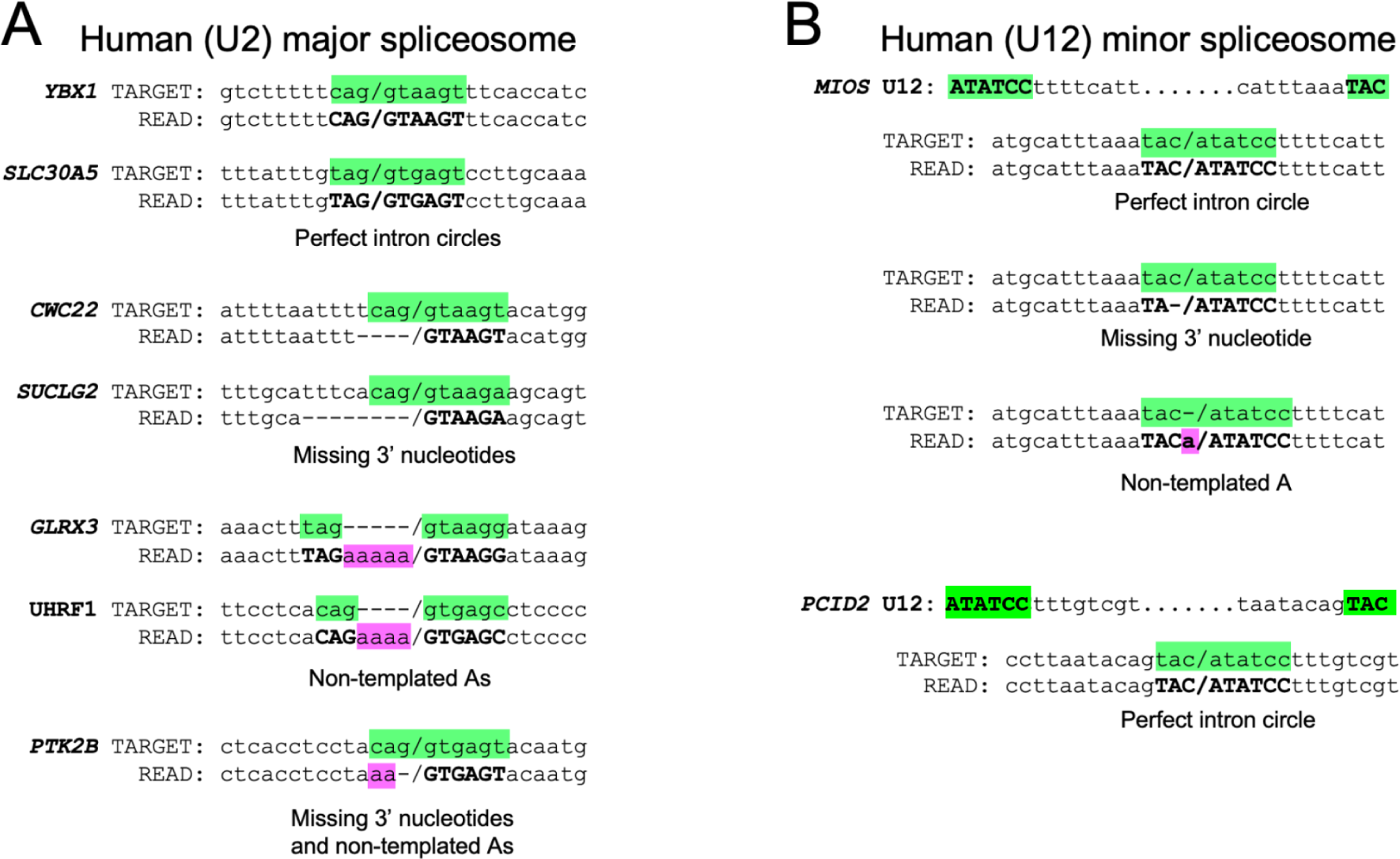
Evidence for full length and processed circles from human cells. (**A**) Selected reads from major (U2) spliceosomal introns found by remapping the branch point enrichment libraries from Mercer et al. (Mercer et al. 2015) to permuted intron target sequences from the human genome. Junctions between the 5’ and 3’ intron ends are denoted by a forward slash. Splice site nucleotides from the permuted target intron are highlighted in green. Dashes are added to the target sequence to maintain alignment when non-genomic As are present in the read. Nucleotides in the read that match the splice sites are in bold, non-genomic As are highlighted in pink. Dashes are added to the read sequence to maintain alignment where nucleotides from the intron 3’ end are missing. (**B**) Selected reads from minor (U12) spliceosomal introns found by remapping the branch point enrichment libraries (Mercer et al. 2015) to permuted intron target sequences from the human genome. Highlight and dash placement are the same as for (**A**). Additional examples with read identifiers are shown in Table S6.

## DISCUSSION

### A model for formation of circular intron RNAs by a catalytically active ILS

Our observations support a model that explains the formation of intron circles (Fig 7). Circles possess an intact 5’ss at closing junction whether processed or not (Fig 1), and are resistant to Dbr1 (Fig 3) as would be expected if the 3’-OH of the lariat tail attacks the branch phosphate to form a 3’-5’ linkage (Reaction F3, Fig 7A, B). The origin of the processed circles is explained by lariat tails longer than about 17 nucleotides visiting the nuclear 3’ processing machinery outside spliceosome before returning to the E1 binding site (Fig 5, see Fig 7C). The E1 binding site is known to accommodate a wide variety of exon sequences during forward splicing, consistent with the many different circle junction sequences observed (Fig 1C, Fig S1). Circles are associated with the spliceosome and Trl1, the only other RNA ligase in yeast, is not required (Fig 2), indicating that this reaction is catalyzed by the spliceosome. The cartoon in Fig 7C shows an overall pathway for formation of the observed classes of intron circles. Since the same classes of circles arise from introns of the major and minor spliceosome (Fig 6), this mechanism appears general for all spliceosomes and most introns across eukaryotes.

**Figure 7.**
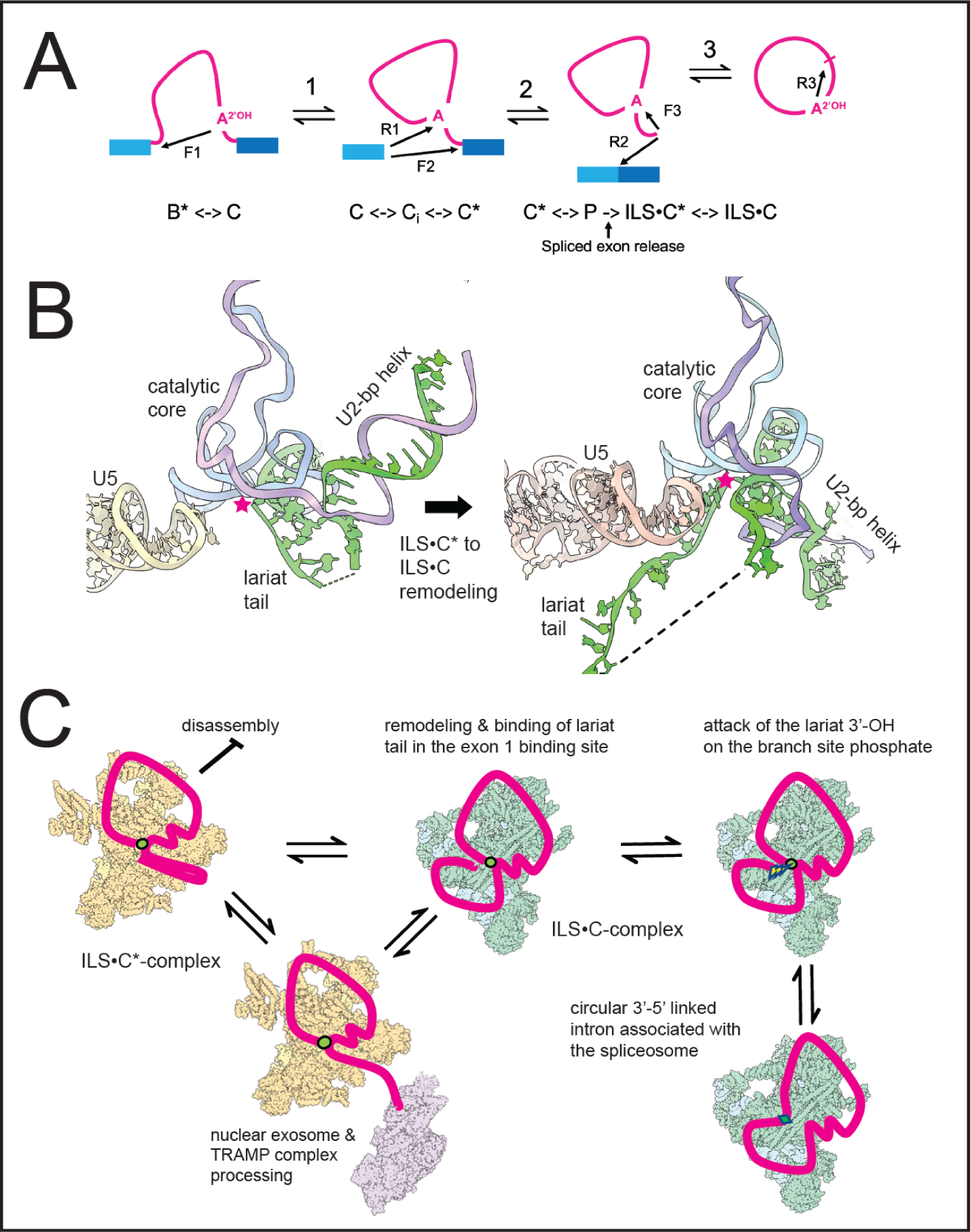
Model for the formation of intron circles. (**A**) Transesterification reactions catalyzed by the spliceosome. F, forward reaction, R, reverse reaction, numbered 1 through 3. The arrows indicate the direction of nucleophilic attack. F1: Attack of the bp 2’-OH on the phosphate (P) at the 5’ ss. F2: Attack of the 3’-OH at the end of E1 on the P at the 3’ ss. F3: attack of the lariat tail 3’-OH on the BP P to form the intron circle. Below the pathway are spliceosome complexes within which these reactions occur. The active site of the C-complex configuration has the 5’ss P and the bp 2’-OH in the active site for F1, whereas the C*-complex configuration has the U2 BP helix pulled out of the active site to create the 3’ss binding site for F2. ILS, intron lariat spliceosome. ILS is formed by spliced exon release (arrow), in the C* configuration (ILS•C*). For F3 it must remodel to the C configuration (ILS•C). (**B**) Cartoon showing remodeling of the RNA in the core of the ILS•C* complex to the proposed ILS•C complex and docking of the lariat tail for circularization. (**C**) Cartoon showing events leading to mixtures of processed and unprocessed full length intron circles. Yellow spliceosomes represent ILS•C*-complex, green represent ILS•C-complexes. The nuclear exosome (purple) represents nuclear 3’ processing machinery interacting with ILS•C* but it is not known which ILS forms permit processing. Intron is in red, key linkages are: green circles: 2’-5’ linkages; green diamonds: 3’-5’ linkages. Lightning bolt indicates nucleophilic attack. Cartoons were made using ChimeraX as above.

The spliceosome is already known to catalyze a reaction similar to the circularization reaction, as revealed by the discovery of spliceosomal reversibility (Tseng and Cheng 2008, 2013). The first forward step of splicing (F1, attack of the bpA 2’-OH on the 5’ss P, Fig 7A) is readily reversible *in vitro* (R1, attack of the E1 3’-OH on the branch P), to recreate the pre-mRNA (Tseng and Cheng 2008, 2013). Our proposed forward circularization reaction F3 is formally similar to R1, replacing the lariat tail for E1 (Fig 7B, left, star). After exons are released, the lariat tail would leave its position in the intron lariat spliceosome (ILS) and find its way to the E1 binding site. During forward splicing, the active site is remodeled from a C-complex conformation (Wan et al. 2016; Galej et al. 2016; Bertram et al. 2020) to C* (Yan et al. 2017; Fica et al. 2017; Bertram et al. 2017) on its way to completing exon ligation in the P-complex (Liu et al. 2017; Wilkinson et al. 2017; Zhan et al. 2022). Our model necessitates that the ILS, which retains the C* configuration (Wan et al. 2017; Zhang et al. 2019; Yan et al. 2015), must return from the C* configuration (ILS•C*) to a C configuration with the branch P in the catalytic center (ILS•C). Once the lariat tail docks in the E1 binding site of ILS•C (Fig 7B, right, star), the reaction can proceed (Fig 7B). The requirement for this remodeling seems a fair expectation given that P-complex spliceosomes (still carrying spliced exons) must also remodel backwards from C* to C while retaining catalytic activity (R2 and R1, Fig 7A) during reverse splicing (Tseng and Cheng 2008). The ILS•C* complex would need to perform a parallel set of rearrangements (Fig 7A), but in the absence of bound exons. Recent refinements of C-complex-like spliceosomes identified the C_i_ complex, a low free energy state complex on pathway between C and C* (Wilkinson et al. 2021; Aupič et al. 2023), further demonstrating the flexibility of spliceosome transitions between the C and C* complexes. This study also identified a long suspected monovalent metal binding site (Tseng and Cheng 2008; Marcia and Pyle 2012; Tseng and Cheng 2013) that when occupied by K^+^, favors F1 over R1 during the first step (Wilkinson et al. 2021; Aupič et al. 2023). This predicts that the proposed circularization reaction (F3), like R1, may be favored when K^+^ is absent or if Li+ is bound in the monovalent metal site of the catalytic core.

### Biological implications of a persistent and active ILS

There are two consequences of extending the lifetime of the ILS: (1) slowing of overall splicing rate, and (2) persistence of a catalytically active spliceosome with empty exon binding sites. Slowing spliceosome disassembly quickly inhibits the early steps of splicing (Mendoza-Ochoa et al. 2019), suggesting that rapid disassembly of the ILS helps satisfy the demand for continued splicing. The consequences of continued catalytic activity of the ILS are less clear. The circular RNAs described here are rare (Fig 1) and in most cases likely harmless, but an active ILS might also catalyze events we have yet to detect *in vivo*. Unusual full length linear forms of introns have been described (Morgan et al. 2019; Yao et al. 2022), and in yeast have been implicated in growth control (Parenteau et al. 2019; Morgan et al. 2019), but whether these are formed within the ILS is not yet clear. In addition, circle formation does not conform to the established tail length rule for the much more abundant linear introns described previously (Morgan et al. 2019). Among the most interesting possible activities of the ILS are reverse splicing reactions that might lead to intron transposition, analogous to those of the group II introns.

The importance of controlling the fate of the ILS is underscored by the evolution of robust disassembly processes that recognize and act on the ILS. This control may have evolved later in the evolution of the splicing machinery, because different organisms seem to have arrived at somewhat different solutions. The structures of the ILS from human, *S. pombe*, and *S. cerevisiae* and their genomes suggest that overlapping but divergent sets of proteins to block catalysis and promote Prp43 binding for disassembly (Raut et al. 2019; Zhang et al. 2019). Among these is the cwf19 protein from *S. pombe* and its human homolog CWF19L2, which carry a conserved domain that binds to and blocks the catalytic center (Yan et al. 2015; Zhang et al. 2019). *S. cerevisiae* has Drn1, a distant cwf19 homolog that promotes Dbr1 activity (Garrey et al. 2014), but Drn1 is absent from the *S. cerevisiae* ILS structure model (but see below). Human and *S. pombe* homologs distantly related to *S. cerevisiae* Spp382/Ntr1 are not found in those ILS complexes, and Ntr2, an essential binding partner of Spp382/Ntr1, has no significant homolog in the *S. pombe* or human genome. The J-protein Cwc23 interacts with and supports the splicing function of Spp382/Ntr1, Ntr2, and Prp43 (Pandit et al. 2009; Su et al. 2018; Sahi et al. 2010), but appears to have divergent functions in other organisms (Raut et al. 2019). The protein compositions of ILS structure models from different organisms may differ due to varying stabilities and abundances of different states of the ILS found in each, and may not be strictly comparable. Nonetheless, these differences hint that control of the ILS after spliced exon release could differ in some eukaryotes in a way that might permit a small fraction of ILS complexes to participate in reverse splicing and intron transposition.

### Extending the parallels between group II introns and the spliceosome

Ideas that explain the birth and persistence of introns and the spliceosome in eukaryotic genomes commonly invoke an ancient self-splicing transposable group II intron that spreads itself as (or before) it fragments during its evolution into the modern spliceosome (Lambowitz and Belfort 2015; Lynch and Richardson 2002). This hypothesis has become ever more cemented by the near superimposability of high resolution structures for the spliceosome and group II introns at various steps in forward splicing (Haack et al. 2024; Xu et al. 2023). If group II introns and the spliceosome shared a common ancestor, they each must have retained, acquired, or lost different abilities as their evolution proceeded along divergent paths. The modern group II introns are highly selected for transposition after splicing, but only to new DNA sites that satisfy the sequence requirements for forward splicing, enforced by the same (in reverse) interactions during reverse splicing into DNA (Lambowitz and Belfort 2015). In contrast the modern spliceosome is not known to reverse splice into DNA. As introns became fixed in eukaryotic genomes and the spliceosome became essential for gene expression, it may have lost the luxury of lingering with its intron on the chance that transposition might occur, possibly through the evolution of the rapid disassembly processes discussed above. Although the specific circularization reaction described here may not be on the direct path to transposition, the existence of ILS catalytic activity opens the possibility that the ILS could reverse splice its lariat intron into a new genomic location, creating a new intron.

Structures of group II introns poised or engaged with DNA substrates are revealing how reverse splicing into DNA leads to transposition (Haack et al. 2019; Chung et al. 2022). In particular, comparisons of structures at different steps in reverse splicing document conformational changes in the positions of the branch helix and the catalytic core (Haack et al. 2019) analogous to the transition of the ILS•C* to ILS•C proposed for circle formation, or for reverse splicing into a new location by the spliceosome (Fig 7, see below). A pathway by which the ILS could acquire a reverse splicing substrate for transposition is revealed by cryoEM structures of the ILS (Zhang et al. 2019). After the spliced exons and several proteins are released from the P-complex to form the ILS, conserved conformational changes in several Prp8 domains create an exon release tunnel between the catalytic center and the outside of the spliceosome (Zhang et al. 2019). This tunnel might also provide a path for a single stranded DNA or RNA to access the exon binding sites and the catalytic core of a still-active ILS for reverse splicing. If the modern spliceosome still possesses an ancestral ability to reverse splice into DNA, it has been obscured in most modern organisms, likely because they have so many introns in their genomes that rapid ILS recycling is essential for continued splicing.

### How the genome got its introns: An unfinished story

The modern spliceosome might still be creating new introns. Diverse microbial genomes show clear evidence of recent repeated insertion of sequence families of spliceosomal introns called "introners" (Gozashti et al. 2022; Worden et al. 2009; Simmons et al. 2015). A major class has signatures of protein transposase-mediated spread (terminal inverted repeats, TIRs, and target site duplications, TSDs, LTRs), however a distinct class of introners lack retroposon or DNA transposon features (Simmons et al. 2015; Collemare et al. 2015; van der Burgt et al. 2012). Members of this class are more often found in algal and fungal genomes, are <200 nt long, do not code for proteins, and might be transposed by the spliceosome (Gozashti et al. 2022; Simmons et al. 2015). Using a special reporter to capture new intron insertions in the laboratory, two instances of intron transposition have been observed in *S. cerevisiae* (Lee and Stevens 2016), supporting the idea that the spliceosome could be the catalyst. There are two proposals for how reverse splicing would lead to intron gain. One posits that reverse splicing into an mRNA is followed by reverse transcription and insertion of the intron-containing cDNA into the genome (Lee and Stevens 2016; Roy and Irimia 2009). However, this process also employs the far more abundant spliced mRNA to erase introns, making it unlikely to contribute to net intron gain (Roy and Irimia 2009). We favor the idea that the spliceosome might reverse splice introns directly into the displaced single stranded DNA of an R-loop (Simmons et al. 2015), which would also place the new intron in the correct orientation on the transcribed strand for splicing. Introns tend to suppress R-loop formation (Bonnet et al. 2017), offering the possibility that new intron insertion at certain locations might be adaptive. Often when introners appear, multiple copies of the same intron are repeatedly transposed. The idiosyncratic ability of some introns to form circles more efficiently than others (Fig 1) suggests that variations in intron sequence and structure might predispose certain introns to evade disassembly more efficiently, or to have enhanced rates of reverse splicing. These special features may reveal how introners could access an ancient pathway for intron transposition that may yet live in the modern spliceosome.

## MATERIALS AND METHODS

### Yeast strains, plasmids, and oligonucleotides

*S. cerevisiae* strains deleted for various single genes came from the yeast deletion collection (Giaever et al. 2002). The *spp382-1* mutant strain was provided by Brian Rymond (Pandit et al. 2006). The *TRL1* strains were provided by Jay Hesselberth (Cherry et al. 2018). Jon Staley provided *PRP43* strains and plasmids carrying U6 mutants (Toroney et al. 2019). Details on these resources are presented in Table S7.

### Traditional cloning of circle PCR products

Outward pointing PCR primers were nested with an intron specific RT primer to capture circular intron sequence junctions. RNA was converted to cDNA with SuperScript III using the intron-specific primer in combination with anchored oligo(dT) and random hexamers, and cDNA was purified on Zymogen Clean and Concentrate columns. PCR was done using intron-specific outward pointing primers (Table S7) and Phusion DNA polymerase (NEB). Purified PCR products were incubated with Taq to add a 3’ A residue and cloned using a Topo cloning kit (Invitrogen). Plasmid inserts were sequenced by Sanger methods and the sequence trace and raw read files are in Table S1.

### RNA extraction and synthesis of the circular GFP spike-in control RNA

Yeast cells were grown in YEPD medium at 30°C according to standard protocols (Guthrie and Fink 1991) and harvested in log phase (OD_600_ = 0.3-0.5). About 5 OD_600_ units of cells were extracted by a hot phenol method (Ares 2012) and evaluated using an Agilent Bioanalyzer to assure intactness. Total RNA was used directly for some libraries, for others RNA was depleted of rRNA by the Illumina RiboZero Gold kit for yeast (no longer available), as described in each case below. Additional treatments before library preparation included digestion of the RNA with the enzymes Dbr1 or RNAseR (see below).

Plasmid pFiniteGFP (Perriman and Ares 1998), see Table S7 for the sequence) was cut with Hind III and used as a template for T7 transcription according to the instructions in the Megaprime T7 transcription kit (Invitrogen) overnight. RNA from the reaction was digested with RNAseR, purified and run on 8M urea, 5% acrylamide gels and the very slow migrating circular RNA band was excised and purified. After purification an aliquot was rerun on the same type of gel and inevitably a small amount of 812 nt linear RNA appeared due to breakage during the preparation of the circular RNA. The estimated fraction of broken circles was subtracted from the absorbance measurement of RNA in order to accurately estimate the true number of intact circles in the spike-in sample.

### Dbr1 digestion

Debranching enzyme Dbr1 digestion was performed according to (Qin et al. 2016), with minor modifications. Recombinant Dbr1 (Khalid et al. 2005) at 6 mg/ml in 50 mM Tris-Cl pH 7.0 was a gift from Aiswaria Krishnamohan in Jon Staley’s group. For the gel in Fig 3, RNA oligonucleotides were purchased from IDT. Oligos (20 pmoles) were labeled at their 5’ ends using gamma-^32^P-rATP (3000 Ci/mmole) and polynucleotide kinase (New England Biolabs), and purified on 7M urea, 8% polyacrylamide gels by elution from excised gel pieces. Digests were performed in 20 ul with ∼10^5^ cpm of labeled substrate, 50 ng unlabeled total yeast BY4741 RNA and 0.25–1 ng of enzyme (0.5–2 ul of a 1:10 dilution of a 50 ng/ul (∼1 uM) working stock) in Dbr1 digestion buffer (50 mM Tris–HCl pH 7.0, 4 mM MnCl_2_, 2.5 mM DTT, 25 mM NaCl, 0.01% Triton X-100, 0.1 mM EDTA, and 0.15% glycerol) at 30°C for 10 min. Digestion products were separated on a 20% acrylamide, 8M urea gel in TBE. For library preparation of DBR1-treated RNA, ∼1ug of rRNA-depleted (RiboZero, Illumina, NB: no longer available) RNA from wild type BY4741 or *spp382-1* yeast or ∼3 ug of total RNA from BY4742, *dbr1Δ*::*KanMX4* were digested in a 10 ul reaction with 50 ng of Dbr1 for 10 min at 30°C, and then diluted with 200 ul 0.3M sodium acetate pH 5.2, extracted with phenol:chloroform:isoamyl alcohol (25:24:1), and then ethanol precipitated and resuspended in 20 ul of water. Recovery was >90% and the RNA was used directly for library preparation.

### RNAseR digestion

For library preparation of RNaseR-treated RNA, ∼1ug of rRNA-depleted (RiboZero) RNA from wild type BY4741 or *spp382-1* yeast or ∼3 ug of total RNA from BY4742, *dbr1Δ::KanMX4* were digested in a 10 ul reaction in the reaction buffer provided by the manufacturer (Epicenter Biotechnologies) containing 20 units of RNAseR for 10 min at 37°C, and then diluted with 200 ul 0.3M sodium acetate pH 5.2, extracted with phenol:chloroform:isoamyl alcohol (25:24:1), and then ethanol precipitated and resuspended in 20 ul of water. Recovery was ∼30-40% for the rRNA depleted samples and ∼20% for the total RNA samples. The RNA was used directly for library preparation.

### Library preparation

Five hundred nanograms RNA processed in various ways (see below) were used as input for library preparation. Reverse transcription was performed using SuperScript IV (Thermo) and first strand primers complementary to a region just downstream from the 5’ss of each target intron, preceded by 4 random nt and then preceded by partial Illumina adapters (Table S7). After purification of first strand product on Zymo Clean and Concentrator-5 columns (Zymogen Research), second-strand synthesis was done using Phusion Polymerase (NEB) and a set of primers complementary to the cDNA (i. e. same sense as the RNA) of sequences upstream of the intron branch point, preceded by 4 random nt (UMI) and partial Illumina adapter sequences for the same set of introns. Double stranded cDNA >150 bp was purified using Zymo Select-A-Size columns. Primer pairs for the control GFP spike-in RNA circle were designed similarly using a first strand primer downstream of the group I 3’ss and a second strand primer upstream of the group I 5’ss in the exonic GFP sequences that become circular (Perriman and Ares 1998). The linear control *TUB3* amplicon is conventionally arranged to amplify a similarly sized internal linear segment of the *TUB3* intron (Table S7). Double-stranded cDNA was bar coded and amplified with Platinum Taq Hifi Polymerase (Invitrogen) with two primers that complete Illumina adapter sequences (95°C for 2min, 27 cycles of [95°C for 30s, 65°C for 30s, and 68°C for 2min], and 68 °C for 7 min). Final sequencing libraries were purified by size selecting the PCR product using Ampure XP beads to obtain fragments >250 bp. Details are available in Supplemental Methods.

#### Mapping and quantification of intron circles from amplicon sequencing libraries

Libraries were analyzed as follows (details in Supplemental Methods). All reads were filtered to have UMI bases with quality scores >33 and paired end reads were trimmed to 35X24 bases. Trimmed reads were mapped using Bowtie2 against fasta files for each intron, the linear control, and the cGFP spike-in to capture reads derived from each target element. Reads mapping to each target element were filtered to ensure they have the correct orientation and the correct distance between read1 and read2 and contain the correct string from the 5’ss into the intron (the filter range, "RangeE"). Filtered reads for each are deduplicated and counted by their UMIs and the top 30 unique reads are used to query the target with BLAT to determine the extent of the 3’ tail match. The mapping and filtering details are described in the Supplemental Methods and analysis parameters are reported in Table S8, and the mapping data is reported in Tables S9-S11.

### Detection of intron circles in spliceosomes

To test for association of intron circles with spliceosomes we employed the methods described in (Shi et al. 2021) with modifications. Briefly, we tagged *CEF1* in the *spp382-1* mutant strain using a PCR product with homology arms targeting the C-terminus of the *CEF1* coding region to insert the TAPS tag in which the calmodulin binding domain is replaced with a StrepTag (Schreieck et al. 2014), a kind gift of Dr. Kelly Nyugen). We modified the plasmid by swapping the 1446 bp BglII-EcoRI fragment carrying the *KanMX6* cassette for the 1333 bp BglII-EcoRI *HIS3MX6* cassette carrying the *Lachancea kluyveri HIS3* gene from a plasmid designed for C-terminal GFP tagging (Longtine et al. 1998) to make pFA6a_TAPS_kHISMX6. Integration was confirmed by PCR across both junctions (see Table S7 for oligonucleotide and plasmid sequences). Cultures were grown at 23°C (*spp382-1* is temperature sensitive), washed and flash frozen in liquid nitrogen as described (Shi et al. 2021), and processed in a ball mill at liquid nitrogen temperature. Extract was bound to IgG-Sepharose 6 Fast Flow resin and washed according to Shi et al. (Shi et al. 2021), except that instead of TEV cleavage, the beads were extracted with phenol:chloroform: isoamyl alcohol (25:24:1) and the aqueous phase was ethanol precipitated. Aliquots of RNA from the input extract, the supernatant after bead binding and the beads were adjusted to volume equivalence with respect to the input extract and subjected to primer extension with ^32^P-labeled oligonucleotides complementary to U2 and SCR1 (Table S7) according to (Perriman and Ares 2007), shown in Fig S2, or were subjected to RT-PCR as described above. The *ECM33* primer pair produces a 227 bp product on the full length circle from the *ECM33* intron as shown in Fig 2.

### Identification of reads derived from circular intron RNAs in published RNAseq datasets

A target fasta database was constructed using all annotated introns in the human genome (hg19), obtained from the UCSC genome browser using UCSC "Known Genes" and the refSeq annotation tracks (Raney et al. 2024). Individual target files were made by joining the 50 bases from the 3’ end of the intron upstream of the 50 bases from the 5’ end of the intron, creating a 100 bp target with a junction similar to the full intron circle in the middle, for each intron. Each target in this initial database was used to query the genome by BLAT and any targets with more than 3 hits were removed.

Sequencing reads from Mercer et al. (Mercer et al. 2015), samples SRR1049823 and SRR1049825 were converted to fasta and used to query the refined target database by BLAT. The resulting alignments were further filtered leaving only the longest target alignment and longest read alignment. Hits to the same target are grouped together in the output file for review. To identify yeast reads from intron circles we followed a similar process using annotated splice junctions from the track "Talkish Standard Introns" on our yeast genome browser at intron.ucsc.edu (Talkish et al. 2019) and the sequencing data in GEO at GSE90105 from (Talkish et al. 2019) and from (Hunter et al. 2024) in the SRA at PRJNA972189. A github deposit with scripts and more technical details can be found at: https://github.com/donoyoyo/intron_circle_hunt

### Data deposition

Sequence data from amplicon pools for all libraries is available under BioProject Accession PRJNA739208 at the Sequence Read Archive https://www.ncbi.nlm.nih.gov/sra

## Supporting information

Zipfile_ALL_suppl_Tables&Text

## ACKNOWLEDGEMENTS

Funding R01 GM040478 and R35 GM145266 to MA. Thanks to Chris Vollmers for advice on library design, Aiswarya Krishnamohan (Staley lab) for Dbr1, Brian Rymond, Rebecca Toroney, Klaus Nielsen, Jon Staley and Jay Hesselberth for strains, Kelly Nguyen for the TAPS plasmid, and Anna Marie Pyle, Anita Hopper, Jon Staley, Jay Hesselberth, Nav Toor, Alex Worden, Jen Quick-Cleveland, Sebastian Fica, Soo-Chen Cheng, and Russ Corbett-Detig for enthusiastic discussions and comments. Special thanks to Doug Black, Tracy Johnson, and Peter and Olivia Narins for providing amazing sabbatical support of different kinds.

## Supplemental Figures and Methods

### Supplemental Figures

**Figure S1.**
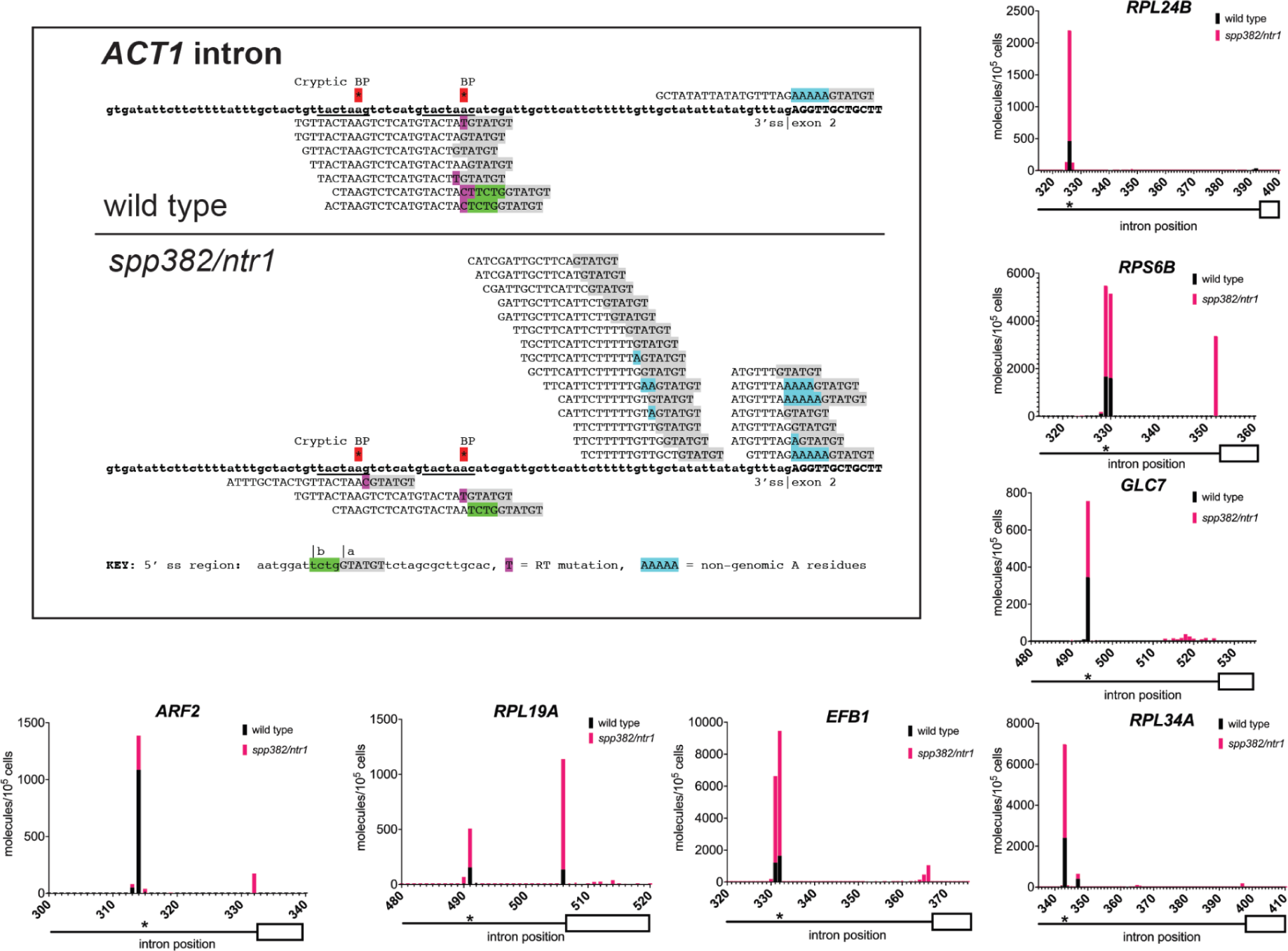
Relevant to Figure 1. Detection of intron circles for additional introns of *S. cerevisiae*. Upper left panel shows circular intron amplicon reads mapping to the *ACT1* intron from wild type (above the line) and the *spp382-1* mutant (below the line). The sequence of the intron is shown in bold. Circle reads are aligned above and lariat reads below the intron sequence with the 5’ss sequence highlighted in gray and non-genomic As highlighted in blue. A class of lariats derived from incorrect 5’ss selection are shown in green. RT errors at the branch are highlighted in purple, see key at bottom. The small graphs show distinct distributions of processed and unprocessed intron circles from different introns. Graphs show the junction locations (x-axis) and number of unique reads with junctions at that location (y-axis, calibrated to a spiked-in circular RNA) per 10^5^ yeast cells, from wild type (black bars) or *spp382-1* (pink bars) cells. The 3’ part of each intron (line) and its second exon (white box) are shown below the x-axis, with the asterisk indicating the position of the bp.

**Figure S2.**
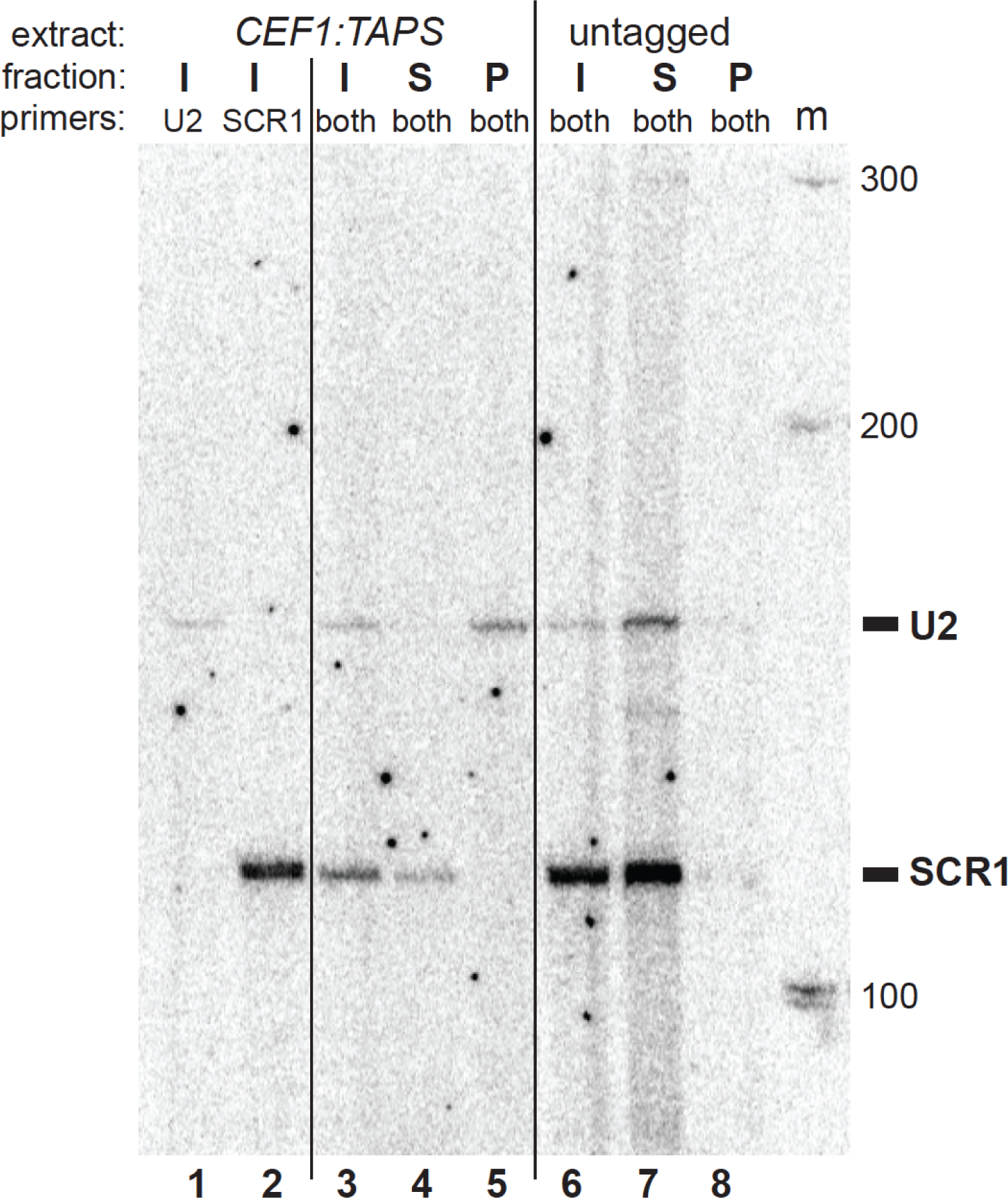
Relevant to Figure 2. Partial purification of spliceosomes by IgG-sepharose chromatography of a CEF1-tagged extract. The *spp382-1* mutant was fitted with a TAPS tag at the C-terminus of its *CEF1* coding region for affinity purification of spliceosomes. Extracts from this strain and an untagged control were prepared (Input, I) and bound to IgG-sepharose. Unbound (Supernatant, S) and bound (Pellet, P) fractions were prepared. RNA from each fraction was used as a template for primer extension with a ^32^P-labeled oligonucleotide complementary to U2 snRNA (lanes 1, 3-8) and SCR1 (lanes 2-8). Labels on the right indicate the expected migration of the cDNA from each primer, 152 nt U2 cDNA primed by oligo 23T, and a 113 nt SCR1 cDNA primed by oligo SCR1-110 (Table S7). The pellet fraction from the tagged extract (lane 5), but not the untagged extract (lane 8) is enriched in the spliceosomal U2 snRNA, free of the cytoplasmic RNA SCR1, which remains in the supernatant fraction of both extracts (lanes 4 and 7).

### Supplemental Tables

Supplemental Table S1: Data for Fig1, Fig S1

Supplemental Table S2: Data for Fig 2C

Supplemental Table S3: Data for Fig 3B, C

Supplemental Table S4: Data for Fig 4

Supplemental Table S5: Data for Fig 5

Supplemental Table S6: Data for Fig 6

Supplemental Table S7: Yeast strains, Plasmids, Oligonucleotides

Supplemental Table S8: Mapping parameters

Supplemental Table S9: Mapping data for Trial 1

Supplemental Table S10: Mapping data for Trial 2

Supplemental Table S11: Mapping data for Trial 3

## Supplemental Methods

### Experimental Trials and Library Details

Trial 1 consisted of 12 libraries designed to test the amplicon method for several introns and to ask about two mutations in spliceosome disassembly proteins (*spp381-1* and *drn1Δ*) and the nuclear 3’ processing machinery (*trf4Δ*, *trf5Δ*, and *rrp6Δ*) using total log phase RNA from wild type and the mutants. The first set of 6 libraries (1.1-1.6) created amplicons for 7 introns (*ACT1*, *EFB1*, *GLC7*, *RPL19A*, *RPL24B*, *RPL34A*, *RPP1B*) in wild type (1.1); *spp382-1* (1.2); *trf4Δ* (1.3); *trf5Δ* (1.4); *rrp6Δ* (1.5); *drn1Δ* (1.6). The second set of libraries (2.1-2.6) created amplicons for 3 introns: *ACT1*, *RPL17B*, *RPL24B* in wild type (2.1); *spp382-1* (2.2); *trf4Δ* (2.3); *trf5Δ* (2.4); *rrp6Δ* (2.5); *drn1Δ* (2.6). The libraries were quantified by qPCR using the Illumina i5 and i7 primers, pooled and sequenced by MA on a MiSeq in Doug Black’s lab at UCLA.

Trial 2 consisted of 12 libraries designed to test the effect of Dbr1 and RNAseR treatment and to replicate experiments with certain mutants, and to assess circles in stationary phase. This trial also included a linear control amplicon from the *TUB3* intron. Both sets of libraries in this trial employed primers for the introns in *ACT1*, *ARF2*, *ECM33*, *EFB1*, *GLC7*, *MPT5*, *RPL19A*, *RPL24B*, *RPL34A*, *RPP1B*, *RPS6B*, and *TUB3*. The first set of 6 libraries (3.1-3.6) used rRNA depleted RNA not treated with enzyme: wild type log phase RNA (3.1); *spp382-1* log phase (3.2); *trf4Δ* log phase (3.3); *rrp6Δ* log phase (3.4); wild type stationary phase (3.5); *spp382-1* stationary phase (3.6). The second set (3.7-3.12) are derived from rRNA depleted log phase RNA treated with enzymes: wild type, RNAseR (3.7); *spp382-1*, RNAseR (3.8); *dbr1Δ*, RNAseR (3.9); wild type, Dbr1 (3.10); *spp382-1*, Dbr1 (3.11); *dbr1Δ*, Dbr1 (3.12). These libraries were sequenced on a NextSeq500 by the staff at the UCSC Paleogenomics Lab.

Trial 3 consisted of 16 libraries designed to quantify the numbers of circles per cell using a spiked in circle, and to extend the test of disassembly factor mutants in altering circle numbers as well as to assess the dependence of circle formation on the tRNA ligase Trl1. RNAs were spiked with an in vitro synthesized 812 nt circular RNA (cGFP) to achieve 1 cGFP molecule per 100 cell equivalents of RNA as follows. Cells were counted at harvest and yields agreed with expected total RNA amounts per cell. A yeast cell contains about 0.7 pg of RNA () so that 10 ug of total RNA arises from 1.43 x 10^7^ cells. At 1 cGFP per 100 cells this requires 1.43 x 10^5^ cGFP molecules. The molecular weight of an 812 nt circular RNA is 2.7 x 10^5^ daltons. Our gel purified cGFP stock was 58.1 ng/ul or 215 nM, or about 1.3 x 10^11^ cGFP molecules/ul. We prepared two 1 ml aliquots of water with 8 ug of HEK293 cell total RNA as a carrier and transferred 1 ul of the cGFP stock (1.3 x 10^11^ molecules) to the first ml, and took 1 ul from that (1.3 x 10^8^ molecules) and diluted it into the second ml of 8 ug/ml HEK cell total RNA to reach a final concentration of 1.3 x 10^5^ cGFP molecules per ul. We then added 1.1 ul of this second dilution (1.43 x 10^5^ cGFP molecules) to each 10 ug of total yeast RNA (from 1.43 x 10^7^ cells) to be used for library creation. All samples were then treated with RNAseR as described below. [Note that comparison of untreated vs RNAseR treated libraries (Trial 2, Fig 3C) shows that RNAseR reduces circular read counts and thus the experiments in Trial 3 may underestimate the true number of circles.] The libraries in this trial employed primers for the introns in *ACT1*, *ARF2*, *ECM33*, *EFB1*, *GLC7*, *MPT5*, *RPL19A*, *RPL24B*, *RPL34A*, *RPP1B*, *RPS6B*, *TUB3*, and the synthetic spiked in circle cGFP. The first set (4.1-4.8) consisted of two wild type replicates 1 and 2 (4.1, 4.2); two *spp382-1* replicates 1 and 2 (4.3, 4.4); two *rrp6Δ* replicates 1 and 2 (4.5, 4.6); and two *trf4Δ* replicates 2 and 3 (4.7, 4.8). The second set (4.9-4.14) consisted of pairs of strains, one wild type control and a mutant: *PRP43* wild type control (4.9); *prp43*-Q423N (4.10); U6 wild type control (4.11); U6*Δ*5 (4.12); *TRL1* wild type control (4.13); *trl1Δ* + *Δ*intron-tRNAs plasmid (4.14). The third set replaced Superscript IV with TIGRT in the first library step and compared wild type (4.15); with *dbr1Δ* (4.16). These libraries were sequenced on a MiSeq by the staff at the UCSC Paleogenomics Lab.

Raw data from all libraries is available in the Short Read Archive under the accession number PRJNA739208. Data in Fig 1 and Fig S1 came from Trial 3 libraries 4.1-4.4, except for the *ACT1* data in Fig S1, which came from Trial 1 libraries 2.1 and 2.2. Data from Fig 2C came from Trial 3 libraries 4.13 and 4.14. Data from Fig 3B and C came from Trial 2 libraries 3.2, 3.8, and 3.11. Data from Fig 4 are from Trial 3 libraries 4.1, 4.4, and 4.9-4.12. Fig 5 data comes from Trial 3 libraries 4.1-4.8.

### Detailed description of circular intron amplicon read processing

The following process was applied separately to each of the samples. Table S8 lists the parameters specific to each sample.

### STEP1: UMI QUALITY FILTERING

The paired-end fastq files were first filtered for high Illumina quality value in all of the positions of the sequenced reads corresponding to the unique molecular index (UMI). This included 4 bases in read1 and 4 bases in read2. The minimum quality value (minQV) was chosen based on the distribution of quality values in those bases, which is dependent on the sequencing platform.

The most common quality value was consistent in those bases for each sample and was above minQV. This step was taken to ensure that when duplicate reads with the same UMIs were removed in a later step, the UMI bases were accurate. This minimizes the possibility that a true molecular duplicate would appear to have a different UMI and thus be erroneously retained. All subsequent steps are performed using these minQV fastq files.

### STEP2A: BOWTIE MAPPING FILTERING

For each intron and the circular cGFP spike-in, the paired-end fastq files were filtered to create a set of intron-specific fastq files using a procedure based on bowtie mapping as follows. A fasta file was created using the full length intronic sequences from the sacCer3 assembly (see Table S8). Each sequence corresponds to the rna-coding strand beginning at the 5’ splice site (GT…) and ending at the 3’ splice site (…AG). A bowtie 2 (Langmead and Salzberg 2012) target was created using this fasta file. For the purposes of mapping, the paired-end 76×76bp minQV fastq files were trimmed to paired-end 35×24bp by removing the first 4 UMI bases and the last 37 or 48 bases from read1 or read2, respectively. These trimmed fastq files were mapped to the bowtie2 target using the bowtie2 parameters "--end-to-end --sensitive --fr" (all others default values). The intent of this mapping is to filter the sequenced fragments to those having the expected primers and subsequent target bases in the introns (or cGFP spike-in). Because the primers are not in the standard orientation for bowtie mapping, and to eliminate spurious hybridization products, the bowtie mappings were further filtered to generate a set of read-IDs as follows. For the circular intron (and cGFP) targets, the bowtie SAM-format output was filtered with samtools (https://github.com/samtools/hts-specs) and the unix tool "awk", based on the expected strand of the read1/read2 primers, and the expected distance between them in the intron.

To extract the read1 mapping IDs for one intron (a "reference" in SAM nomenclature) this sequence of commands was used:

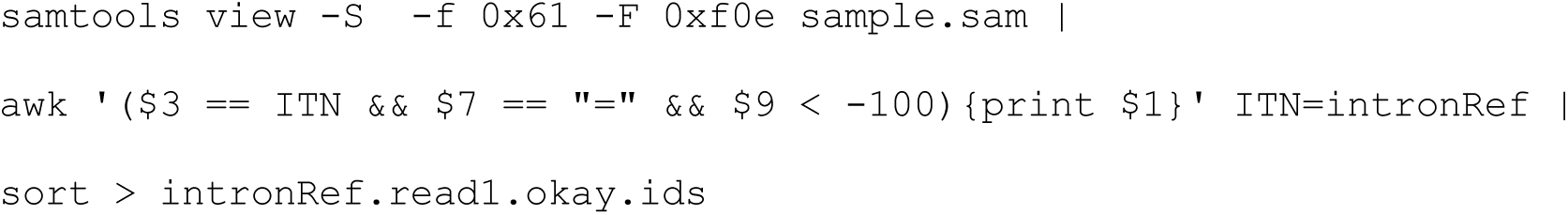

To extract the read2 mapping IDs:

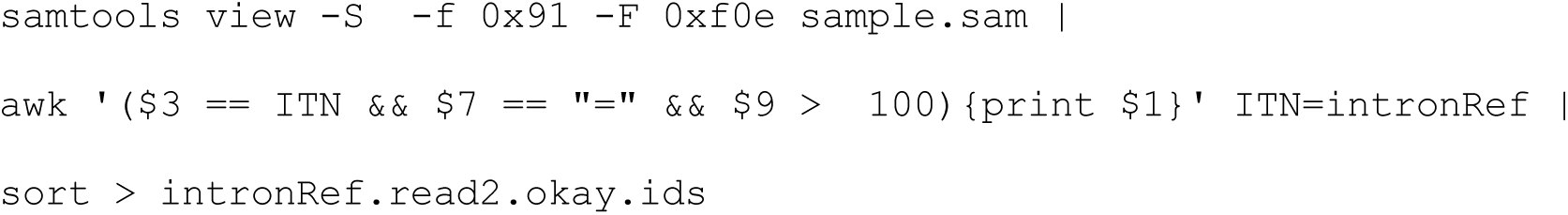

The "-f" and "-F" options together specify: read1 or read2, its strand, the fact that both reads of the pair are mapped, and that this is NOT a "proper pair" by the standard bowtie2 determination. The third field ($3) of the SAM file is selected for the particular reference, and the seventh field ($7) specifies that the other read maps to the same reference. The sign and magnitude of the ninth field ($9) indicates the minimum required distance between the read locations in the intron as well as their relative positions in the intron. The first field ($1) is the desired read-ID. The 2 files of read-IDs were verified as identical, and either can be used in the next step. For the non-circular *TUB3* intron target, slightly different parameters were used because of the expected position and orientation of its primers.

For read1 of the linear *TUB3* intron target:

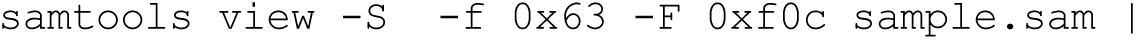
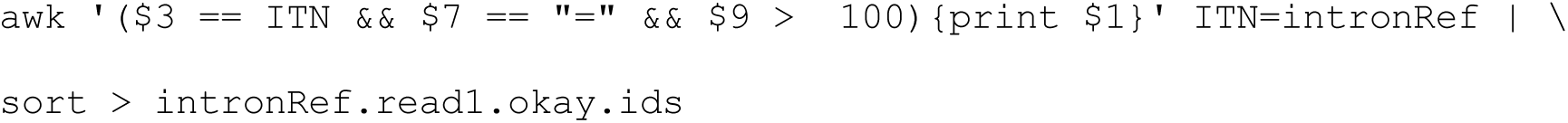

For read2 of the linear *TUB3* intron target:

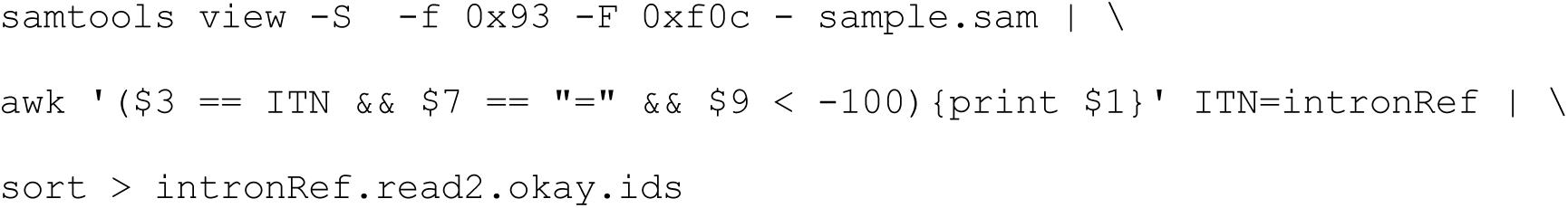

Using the set of read-IDs for a given intron (whether circular or linear target), the original full-length paired-end fastq files were filtered to create a set of intron-specific fastq files. All subsequent steps are performed using these intron-specific fastq files.

### STEP2B: EXPECTED SEQUENCE FILTERING

As indicated in Table S8, a specific range of bases from read2 in the fastq files (filtered as described in the previous step) were extracted. Only those fastq entries that had an exact match to the intron-specific expected sequence string were retained for the following analysis step. The range and expected string for this step were chosen based on the intronic bases adjacent to the 5’ splice site of the intron (or the expected circularization point of the cGFP spike-in).

### STEP3: SEQUENCE ANALYSIS INCLUDING DUPLICATE REMOVAL

As indicated in Table S8, a specific range of bases from read2 were analyzed for each intron. This range was chosen based on the expected position in read2 of the 5’ splice site (5’ss) of the intron, relative to one of the primers used to extract the RNA fragment sequenced. In particular, the analysis bases were chosen to overlap the 5’ss position and include 6 expected intronic bases upstream and 14 unknown bases from the downstream part of the intron. The downstream bases should differ depending on whether the sequenced fragment was from a lariat, an RNA circle, or something else. The upstream bases might also differ from the expected string if the spliceosome used an incorrect 5’ss upstream of the expected 5’ss, but incorrect downstream 5’ss usage would be filtered out in the previous sequencing filtering step. To remove duplicates from the set of analysis strings, the following algorithm was used. First, the analysis string was extracted for all paired-end reads into a file "analysis.all". Similarly, the UMI bases from read1 and read2 were extracted into files "umird1.all", "umird2.all". To collapse duplicate strings having the same UMI, the following unix string of piped commands was executed:

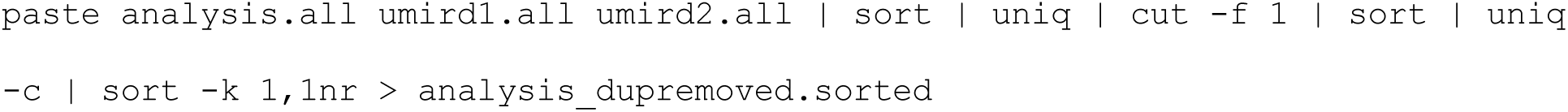

The "paste" command associates each occurrence of a string with its UMI. The initial "sort | uniq" command removes all duplicates, leaving only one copy of a string-umi combination. The "cut" command strips off the UMI, leaving the dup-removed analysis strings. The next "sort | uniq -c" command counts occurrences of each distinct dup-removed string. The final "sort" command arranges these in decreasing order of counts. These counts and their sum are combined to generate the fraction and cumulative fraction for these distinct dup-removed strings, which can also be presented in reverse-complement form for easier downstream analysis. Each string represents a distinct 5’ss-3’ intron end junction including the 6 bases of the 5’ss, and the counts of each represent their abundance in the sample.

### STEP 4: MAPPING THE JUNCTION ENDS OF EACH UNIQUE CIRCLE

To identify the circle junction, we used BLAT (Kent 2002). The last 6 nt of the string were removed and each remaining sequence was used to query the intron sequences (except GLC7, which was done separately due to low information content segments of its intron). BLAT options used were:

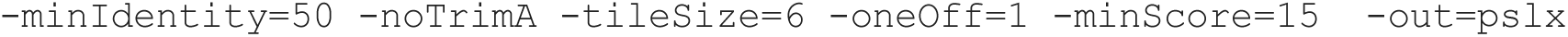

From the pslx file output, the BLAT score (number of matching nucleotides), the position in the intron sequence of the end of the match (this is the nucleotide joined to the 5’ss), and the match sequence were reported along with the query, its number of counts and its rank. These values are reported in the Supplemental Tables S9-S11. Classification and counting of junctions as lariats or circles was done using the nucleotide positions specific for each intron as described in Table S1.

